# Sequence-directed RNA remodeling within a topologically complex RNP substrate

**DOI:** 10.1101/2022.02.24.481786

**Authors:** Victor Emmanuel Cruz, Kamil Sekulski, Nagesh Peddada, Carolin Sailer, Sahana Balasubramanian, Christine S. Weirich, Florian Stengel, Jan P. Erzberger

## Abstract

DEAD-box ATPases are ubiquitous enzymes essential in all aspects of RNA biology. However, the limited *in vitro* catalytic activities described for these enzymes is at odds with their complex cellular roles, most notably in driving large-scale RNA remodeling steps during the assembly of ribonucleoproteins (RNPs). We describe cryo-EM structures of 60S ribosomal biogenesis intermediates that reveal how context-specific RNA unwinding by the DEAD-box ATPase Spb4 results in extensive, sequence-directed remodeling of rRNA secondary structure. Multiple *cis* and *trans* interactions stabilize a post-catalytic, high-energy intermediate that drives the organization of the root helix structure within rRNA domain IV. This mechanism explains how limited strand separation by DEAD-box ATPases is leveraged to provide non-equilibrium directionality and ensure efficient and accurate RNP assembly.

---

DEAD-box ATPases are ubiquitous and highly conserved enzymes involved in all aspects of RNA metabolism, including mRNA processing, export, translation and decay as well as the assembly of multiple cellular RNPs (*1*-*3*). Unlike other superfamily 2 (SF2) helicases, DEAD-box proteins are not processive (*4*) and the structural and biochemical characterization of their biological function has been hampered by the complexity of their RNA and RNP substrates. In addition, most DEAD-box ATPases have very low *in vitro* enzymatic activities. Detailed biochemical studies of model DEAD-box proteins Ded1 and Mss116 revealed that they act by locally unwinding short RNA duplexes to separate up to ∼16 nucleotides of duplex RNA (*5, 6*). Strand separation is achieved using a two-step mechanism (*7, 8*). First, in separate events, the N-terminal D1 RecA domain engages ATP and the C-terminal D2 RecA domain engages dsRNA (Fig. 1A). Second, the two domains cooperatively assemble into a “closed state,” completing ATP and ssRNA binding pockets, stabilizing the D1/D2 interface and disrupting a limited number of base-pairs. This mechanism limits full strand displacement activity to short, topologically unencumbered substrates (Fig. 1A). Product ssRNA engagement is maintained by conserved interactions with the RNA phosphate backbone and is dependent on the presence of ATP; substrate release is triggered by nucleotide hydrolysis(*9*).

**Figure 1.**
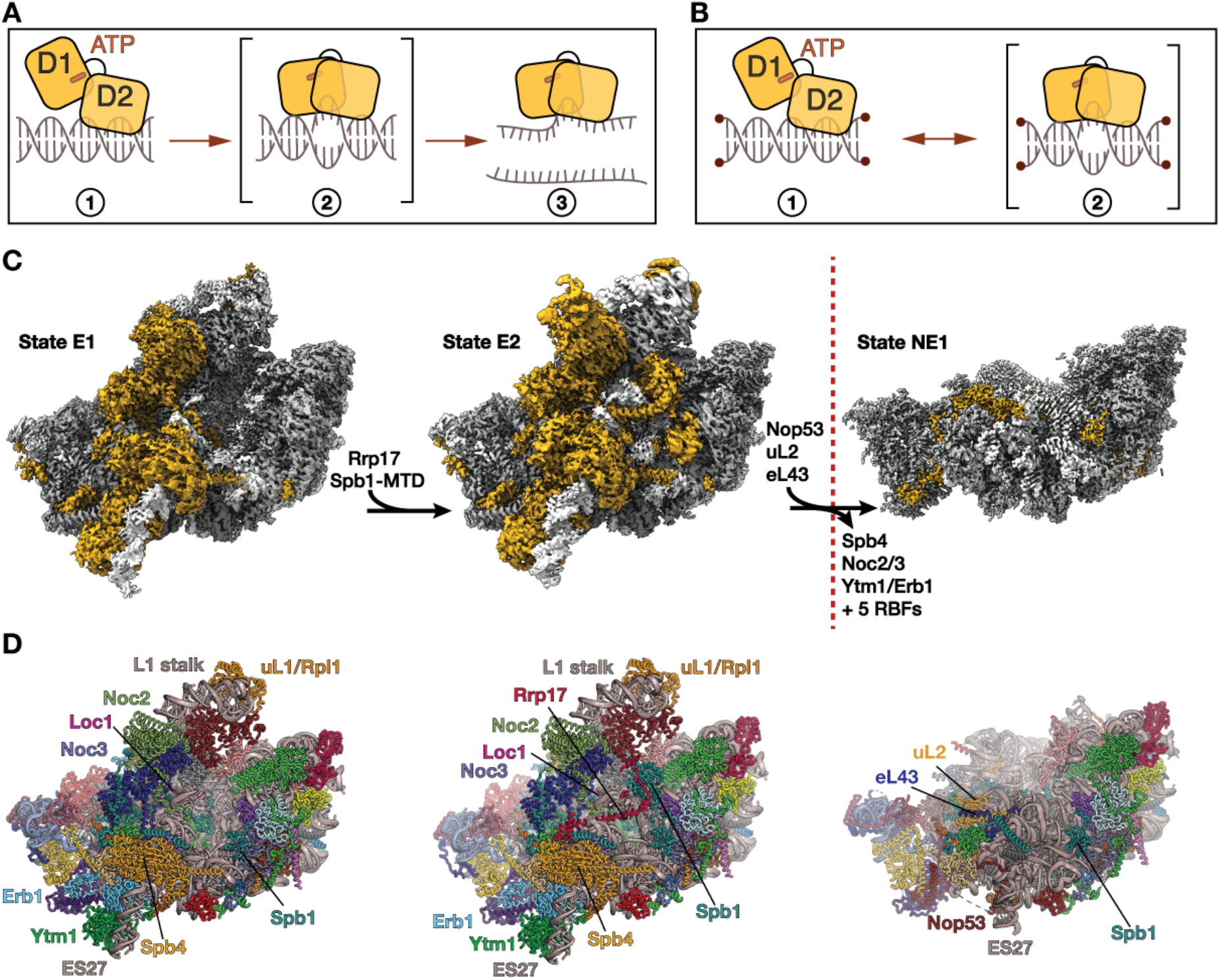
Cryo-EM reconstructions of late nucleolar/early nucleoplasmic pre-60S particles. (A) Schematic of the canonical DEAD-box ATPase dsRNA unwinding mechanism. 1. Core domain D1 is bound to ATP while D2 independently initiates dsRNA engagement. 2. D1 and D2 assemble to unwind a short dsRNA segment, trapping ∼5nt of single-stranded RNA to generate a high-energy intermediate. 3. Expansion of the initial “bubble” within short, unencumbered substrates can result in the full displacement of the complementary RNA strand. (B) Limitations of DEAD-box ATPase activity 1. Initial engagement of ATP and dsRNA by D1 and D2 occurs as in panel A. 2. DEAD-box ATPase binding is unproductive if the substrate is constrained (indicated by the red spheres), as topological strain introduced by local unwinding cannot be dissipated. (C) Cryo-EM maps of late-nucleolar and early nuclear pre-60S particles. rRNA density is shown in white, density for RBFs and RPs interpreted in previous reconstructions are shown in dark gray. Densities of RBFs and RPs with *de novo* or substantially expanded molecular models are colored in gold. The arrows denote maturation progression and listed proteins highlight pre-60S composition changes at each step. The red line indicates the nucleolar/nuclear boundary. (D) Cartoon representation of the modeled structures built into the maps shown in panel C. Polypeptides corresponding to the gold densities in panel C are individually labeled.

This strand separation and release mechanism is consistent with the canonical role of DEAD-box proteins as local disruptors of small duplex RNA elements. Established examples include RNA chaperones that disrupt metastable RNA secondary structures to promote proper RNA folding (*10*), translation initiation modulators that resolve hairpin structures in the 5’ UTRs of mRNAs (*11*), and disassemblers of snoRNA/rRNA duplex structures during rRNA maturation (*12*). However, the majority of eukaryotic DEAD-box ATPases are involved in the assembly of large RNPs (*2, 12, 13*). These complexes contain stable, topologically constrained RNA duplex elements that cannot be readily unwound (Fig. 1B), raising the question of how DEAD-box proteins use limited local strand disruption to remodel thermodynamically stable substrates.

To study DEAD-box activity within a complex RNP, we chose the well-characterized biogenesis of the large ribosomal (60S) subunit. As part of this maturation pathway, the large subunit (25S) rRNA undergoes a series of defined structural transitions, coupled to the ordered incorporation of ribosomal proteins (RPs) (*14, 15*). This mechanism ensures the precise folding and docking of distinct rRNA domains (I-VI) around their root helices (Fig. S1). In the yeast *Saccharomyces cerevisiae*, ∼70 ribosome biogenesis factors (RBFs) guide this process, including six essential members of the DEAD-box family of ATPases: Dbp6, Dbp9, Mak5, Drs1, Dbp10 and Spb4, proposed to have dual functions as RNA remodeling factors and surveillance modules(*16, 17*). Although present in various proteomic and mapping studies of pre-60S intermediates (*17*-*22*), no structural models or defined RNA substrates are available for any of these enzymes. In this study, we focused on Spb4 because, although its molecular structure, precise rRNA substrate, and function in biogenesis were unclear, it had been assigned to a region of weak cryo-EM density within the State E late nucleolar 60S intermediate (*20*-*24*). Cryo-EM reconstructions show that local strand unwinding by Spb4 triggers an extensive, sequence-directed rRNA rearrangement. The stabilization of the resulting high-energy RNP structural intermediate drives the anchoring of the domain IV root helix structure, triggering the docking of this rRNA subregion. Our findings reconcile the limited *in vitro* capabilities of DEAD-box ATPases with the complex *in vivo* activities of these proteins.

## Full atomic models of late nucleolar/early nucleoplasmic pre-60S intermediates

To obtain cryo-EM reconstructions of sufficient quality to characterize the function of Spb4, we enriched Spb4-containing particles *in vivo* by overexpressing a tagged version of RBF Ytm1 that lacks its N-terminal ubiquitin like domain (Ytm1^ΔUBL^) (Table S1 and table S2). In these strains, the large dynein-like AAA+ ATPase Rea1 fails to remove Ytm1^ΔUBL^ from pre-60S particles, halting biogenesis immediately prior to the release of Spb4 and causing the accumulation of nucleolar State E (*25*). Single particle cryo-EM reconstructions of a homogeneous sample obtained by Ytm1^ΔUBL^/Spb4 tandem affinity purification (Fig. S2) generated high-quality maps with overall resolutions of 3Å (Fig. S3, A to K, fig. S4 and table S3). Local 3D classification defined two distinct states, which we call E1 and E2, that represent sequential assembly intermediates (Fig.1, C and D, fig. S4). State E2 is distinguished by the presence of the methyltransferase domain (MTD) of the RBF Spb1 and the newly identified RBF Rrp17 (Fig. S5, A, B and C). Clear density corresponding to Spb4 is present in both E1 and E2 but is of slightly higher quality in state E2. Additional maps were obtained by locally refining five sub-regions centered, respectively, around Spb1, Spb4, the Noc2/Noc3 complex, the L1-stalk/uL1 assembly, and the “foot” structure surrounding internal transcribed sequence 2 (ITS-2) (Fig. S3 and fig. S4). This set of overlapping maps allowed us to interpret and build atomic models into previously unidentified or poorly resolved regions of the State E 60S intermediate. In addition to allowing a full characterization of the Spb4/60S interaction, we built atomic models of the Spb1-MTD/A-loop interaction as well as *de novo* structures for the RBFs Noc2 and Loc1 (Fig. S5, D and E), the RP uL1 and additional structural elements of RBFs Noc3, Cic1, Nop2, Spb1, Ytm1 and Erb1, yielding the most complete and accurate molecular model of the late nucleolar pre-60S intermediate to date (Fig. S3 and table 1). To better understand the immediate structural consequences of Spb4 removal, we also purified 60S intermediates enriched for the RBFs Nop53 and Spb1 via tandem affinity purification from steady state lysates (Fig. S2B, table S1 and table S2) (*24*). A single particle reconstruction of this intermediate to 2.8 Å shows the pre-60S structure immediately after Rea1-mediated Spb4 removal (Fig.1, C and D, fig. S6 and table S3), allowing for a direct structural comparison with the preceding E2 intermediate and revealing a previously uncharacterized helix within the RBF Nop53 that stabilizes expansion segment 27 (ES27) immediately after Spb4 release (Fig. S5F).

## Spb4-mediated strand separation triggers extensive, sequence-specific remodeling of rRNA

A complete atomic model of Spb4 bound to its substrate rRNA on the pre-60S was built into the locally refined maps. The catalytic D1 and D2 domains engage the phosphate backbone of rRNA residues C1941-A1945, which map to rRNA helix 63 (h63) in the mature 60S, in the canonical manner observed in other DEAD-box•RNA complexes (Fig. 2, A and B) (*2*). The h62/h63/ES27 region is dynamic and not resolved in the structures of early nucleolar intermediates, but global SHAPE analysis (*26*) and footprinting studies (*21*) show that the h62/h63/ES27 region forms stable RNA duplexes in the earliest nucleolar pre-60S intermediates and that this base-pairing is disrupted when Spb4 is present (*20*). Consistent with this data, h63 and approximately half of h62 are unwound in the E2 pre-60S particle (Fig. 2C) when compared to their fully paired structures in the NE1 state immediately after Spb4 release (Fig. 2D). While h62 is topologically unencumbered and can be freely unwound, h63 is in a more constrained internal region of the 25S (Fig. 2D and fig. S1). This position disfavors the expansion of any strand separation activity by Spb4, since any resulting reduction in twist cannot easily be compensated by an equivalent change in writhe (i.e. a full rotation of the entire h63/ES27 region). Instead, we find that the local topological disruption caused by Spb4 RNA unwinding is propagated towards the rRNA core, leading to the formation of an alternate RNA helix (h62-alt) involving the bases of h62. The increase in twist upon formation of this compensatory helix maintains the overall RNA topology within h62/h63/ES27, and the formation of alternate base-pairs within h62-alt recovers some of the thermodynamic cost of Spb4-mediated strand separation (Fig. 2C). Thus, h62-alt can only form because of the availability of a complementary sequence in close proximity to the RNA element initially unwound by Spb4 (Fig. 2C), allowing this local dsRNA disruption to induce an extensive, sequence-specific rRNA arrangement. Remarkably, despite the presence of only 5 nucleotides in the active site of Spb4, this mechanism triggers the remodeling of 35 rRNA nucleotides from a single local strand disruption event (Fig. 2, C and D). This mechanism is fundamentally different from the observed functions of DEAD-box ATPases in bacterial and mitochondrial ribosome biogenesis (*27*).

**Figure 2.**
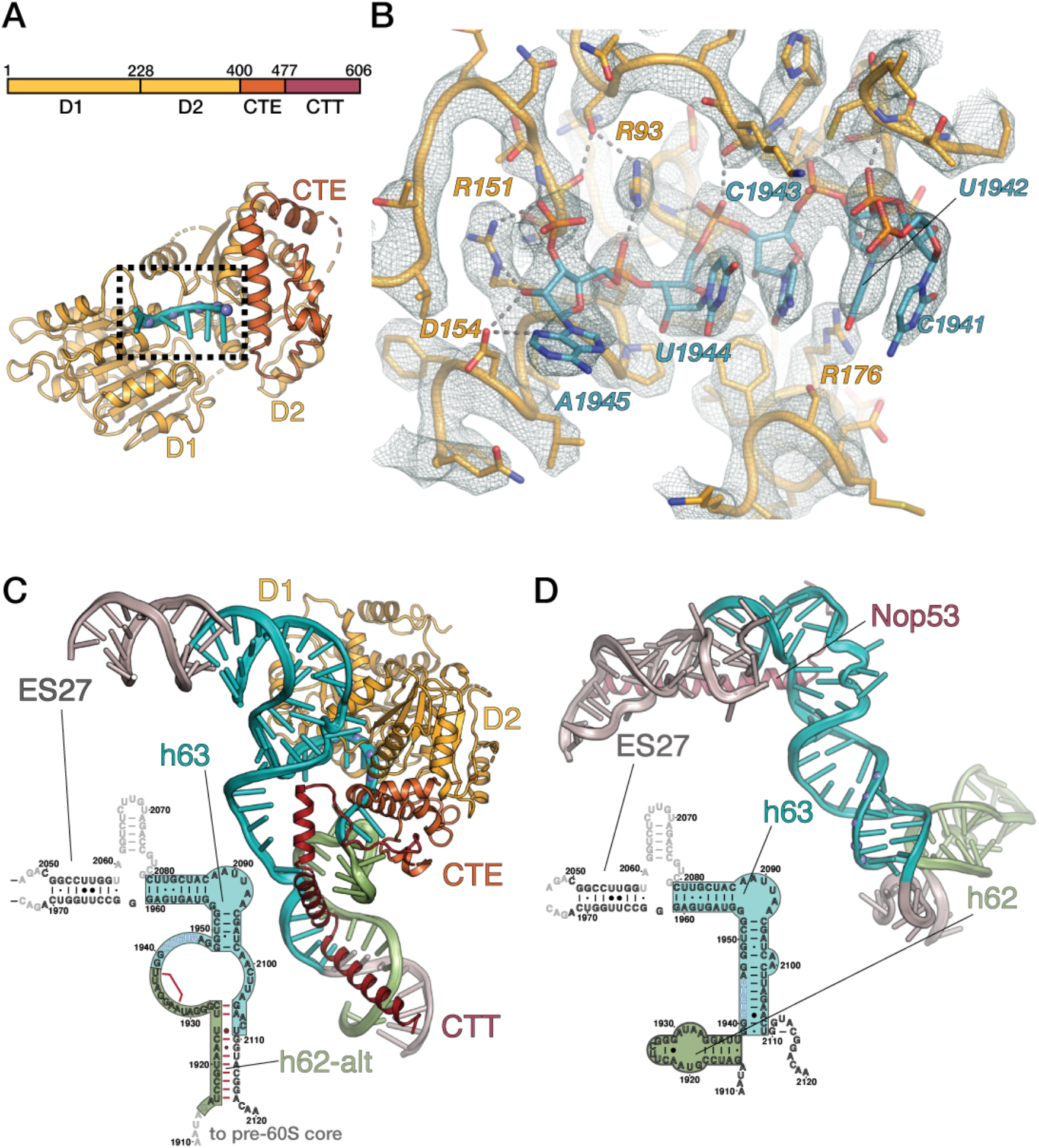
Sequence-directed helix remodeling by Spb4. (A) Top – Domain structure of Spb4. The protein core is composed of dual RecA domains (D1 and D2), a C-terminal extension (CTE) and a C-terminal tail (CTT). Bottom - Cartoon depiction of the D1/D2/CTE Spb4 core showing the five RNA bases bound in the canonical RNA binding pocket at the D1/D2 interface. (B) Detailed view of the active site of Spb4 (corresponding to the boxed region in panel A), engaged to substrate rRNA in state E2. Key Spb4 residues involved in coordinating interactions with the phosphate backbone are shown as sticks and labeled, hydrogen bonding networks are shown as dotted lines. Cryo-EM density of the locally refined map is shown as a mesh. (C) Cartoon representation and base-pairing diagram of the h62/h63/ES27 rRNA region bound by Spb4 in state E2 (colored as in panel A). rRNA bases located in h63 are colored in cyan, those in h62 are colored in green and the equivalent regions are boxed accordingly in the secondary structure diagram. The five residues observed in the active site of Spb4 are marked with blue spheres in the cartoon and shown in white in the secondary structure diagram. The alternate base pairs formed within h62-alt are highlighted in red. (D) Cartoon representation and base-pairing diagram of the h62/h63/ES27 rRNA region as observed in the NE1 structure, immediately following Spb4 release. rRNA regions are highlighted as in panel A and the newly identified central helix of RBF Nop53 is shown in dark red. The five residues observed in the active site of Spb4 in the E2 state are marked with blue spheres in the cartoon and shown in white in the secondary structure diagram.

## *Cis* and *trans* interactions stabilize the remodeled, high-energy Spb4/60S intermediate

The E2 structure therefore represents a post-remodeling RNA “trapped” in a high-energy state stabilized by two *cis* elements within Sbp4 (Figs. 2C, fig. 3, A and B). First, the C-terminal extension (CTE, residues 406-499) packs closely onto the catalytic core of Spb4 and engages the four nucleotides (U1937-G1940) immediately 5’ to those within the D1/D2 RNA binding pocket (Fig. 3A). These rRNA interactions are consistent with CRAC analysis of Spb4 rRNA binding in both yeast and human pre-60S ribosomes (*20, 21*). Unlike the ribose interactions at the D1/D2 interface, the Spb4-CTE interacts with the bases of the rRNA, ensuring sequence specificity in expanding the initial strand separation and promoting the unwinding of h62 to form h62-alt. The second *cis* element engaging the product rRNA is the C-terminal tail of Spb4 (CTT, residues 500-606), which forms a V-shaped α-helical structure that directly engages h62-alt and intercalates a conserved tryptophan residue (W536) at its apex (Fig. 3B).

**Figure 3.**
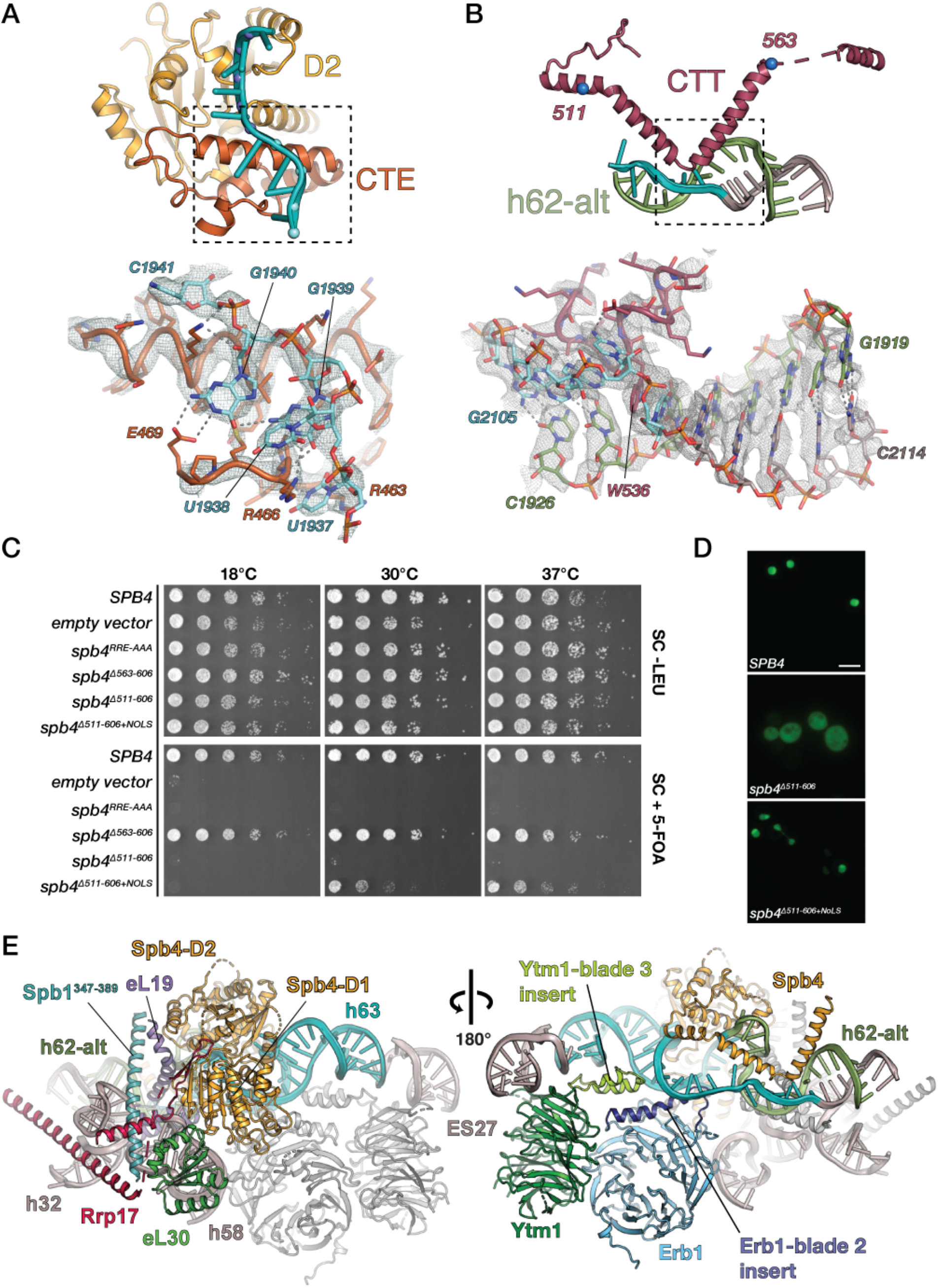
*Cis* and *trans* elements modulate Spb4-directed helix remodeling. (A) Top - Cartoon representations of the h62 bases engaged by the Spb4 D2 and CTE modules, colored as in Figure 2. Bottom – Detailed view of the CTE/rRNA interaction (corresponding to the boxed region) highlighting base specific H-bonds between CTE residues E463, R466 and R469 and individual bases. The cryo-EM density of the locally refined maps is shown as a mesh. (B) Top - Cartoon representation of h62-alt engaged by the Spb4 CTT, colored as in Figure 2. Blue spheres denote the sites of engineered CTT truncations. Bottom - Detailed view of the CTT/h62-alt interaction (corresponding to the boxed region) highlighting the alternate base-pairing within h62-alt and predicted H-bond interactions between CTT residues and individual bases. The cryo-EM density of the locally refined maps is shown as a mesh. (C) Plasmid shuffle assays testing the ability of *SPB4* CTE and CTT mutants to rescue the lethality of *spb4Δ*. Top panel – serial dilutions of cells grown at 18°, 30° and 37° in the presence of p*SPB4*-URA3 and the LEU2 plasmids listed. Bottom panel – serial dilutions of mutant strains after counterselection against pSPB4-URA3 on 5-FOA plates. (D) Fluorescent imaging monitoring the localization of N-terminally GFP tagged Spb4. Spb4 is nuclear/nucleolar, while Spb4^Δ511-606^ becomes cytoplasmic. Reincorporation of a NoLS in Spb4^Δ511-606 + NoLS^ restores nuclear/nucleolar localization. (E)The high-energy intermediate is stabilized by a *trans* interaction network. Cartoon representations viewed from opposite directions. Left – Interaction network involving the RBFs Rrp17, Spb1 and RPs eL19 and eL30 maintain the high-energy state of Spb4. Individual proteins are colored as in Figure 1. Right – conserved β-propeller blade insertions in the RBFs Ytm1 and Erb1 engage and stabilize the junction of the duplex and unwound regions within h63. Proteins are colored as in Figure 1, blade insertion are highlighted in bright green and dark blue and labeled.

The CTE and CTT interactions suggested that both elements facilitate Spb4-mediated rRNA remodeling and are therefore essential *in vivo*. Indeed, the triple mutant *spb4*^*R463A/R466A/E469A*^ (*spb4*^*RRE-AAA*^), designed to disrupt sequence-specific interactions between the CTE and the rRNA (Fig. 3A), is unable to complement the lethality of *spb4Δ* (Fig. 3C). This finding is consistent with our hypothesis that the expansion of the initial ssRNA “bubble” is favored only when h62 is disrupted, allowing formation of h62-alt. *In vivo* characterization of the CTT/rRNA interaction is complicated by the presence of a nucleolar localization signal (NoLS) within this element (*21*). To separate the localization and rRNA-interaction functions of the CTT, we generated three mutant forms of *SPB4*. Deletion of Sbp4 residues 563 - 606 (*spb4*^*Δ563*^), does not affect growth, showing that the most distal elements of the CTT are not essential (Fig. 3C). Expression of a Sbp4 variant truncated at position 511 (*spb4*^*Δ511*^), missing both the h62-alt interaction region and the NoLS, mislocalizes to the cytoplasm and is lethal (Fig. 3C and 3D). Proper cellular localization can be restored with the addition of a heterologous localization signal (*spb4*^*Δ511-NOLS*^) (*28*) (Fig. 3D). Expression of *spb4*^*Δ511-NOLS*^ results in a pronounced temperature-dependent growth defect, ranging from near wild-type growth at 37°C to non-viability at 18°C (Fig. 3C). This cold-sensitive phenotype is consistent with a role for the CTT in facilitating initial strand separation and secondary structure remodeling activities. At higher temperatures, RNA duplexes are less stable, making the CTT non-essential; at lower temperatures, RNA duplex structures are more thermodynamically stable, making the CTT contribution essential.

In addition to the Spb4 CTE and CTT, an extensive network of *trans* interactions composed of RPs, RBFs and rRNA segments helps stabilize the post-catalytic state. Prominent among these are direct interactions with Sbp4; RP eL30, RBF Rrp17 and rRNA h58 interact with domain D1 while RP eL19 and Spb1 engage D2 and the CTE (Fig. 3E – left view). A long helical element within Spb1 (residues 347-389) bridges D1 and D2 to further stabilize the active D1/D2/rRNA interface (Fig. 2A). *Trans* interactions also contribute to post-catalysis stabilization by binding the remodeled rRNA. The β-propeller of Ytm1 binds the first helical segment of ES27, while blade insertions in the β-propeller domains of both Erb1 and Ytm1 stabilize h63 at the junction of the unwound region (Fig. 3E – right view). These insertions are not ordered in the crystal structure of the isolated Erb1/Ytm1 heterodimer (*29, 30*), indicating that they become structured specifically upon engaging the high-energy Spb4 intermediate. Complementing its interactions with the Spb4-CTT, eL19 also interfaces with the h62-alt backbone using the same amino acid residues that mediate backbone interactions with h63 in the NE1 state and the mature ribosome.

## Nucleotide hydrolysis precedes the final stabilization of the Spb4/60S intermediate

Other DEAD-box ATPases have been visualized in a trapped, RNA-bound state, most notably the exon-junction-complex component eIF4AIII (*31*-*33*). Unlike eIF4A-III, whose trapping mechanism within the EJC is nucleotide dependent, Spb4 is stabilized in the absence of bound ATP. The bipartite nucleotide-binding pocket of Spb4 is slightly open, placing key ATP-binding residues beyond hydrogen-bonding range and disfavoring ATP binding (Fig. 4A). Mutations in DEAD-box catalytic motifs I (K57A), II (D172A), required for nucleotide binding, and VI (R360V), required for hydrolysis, fail to complement the lethality of *spb4Δ* (Fig. 4B), indicating that nucleotide binding and hydrolysis are required for Sbp4 function. While the importance of ATP binding for strand separation by Spb4 is clear, the function of hydrolysis in establishing the post-catalysis complex is less intuitive. Overexpression of catalytically inactive Sbp4 variants does not result in a dominant-negative phenotype, unlike overexpression of *ytm1*^*ΔUBL*^ (Fig. 4C). This result is expected for Spb4^K57A^ and Spb4^D172A^, which are unable to effectively engage ATP. However, Spb4^R360V^, which is predicted to bind but not hydrolyze ATP (*34, 35*), is also unable to effectively compete for stable 60S binding, suggesting that ATP hydrolysis is required to establish the stabilized post-catalysis state. Modeling of a closed, ATP-bound conformation of Spb4 indicates that the interaction between D1 and eL30 is more favored in the open state following ATP hydrolysis. To test this idea, we mutated a triad of residues on Spb4 expected to disrupt the eL30 interface (Fig. 4A). The triple mutant *spb4*^*Q196A/R198A/R219A*^ (*spb4*^*QRR-AAA*^) shows a moderate growth defect at 30°C (Fig. 4D), consistent with a role in stabilizing the final, post-hydrolysis conformation of Spb4.

**Figure 4.**
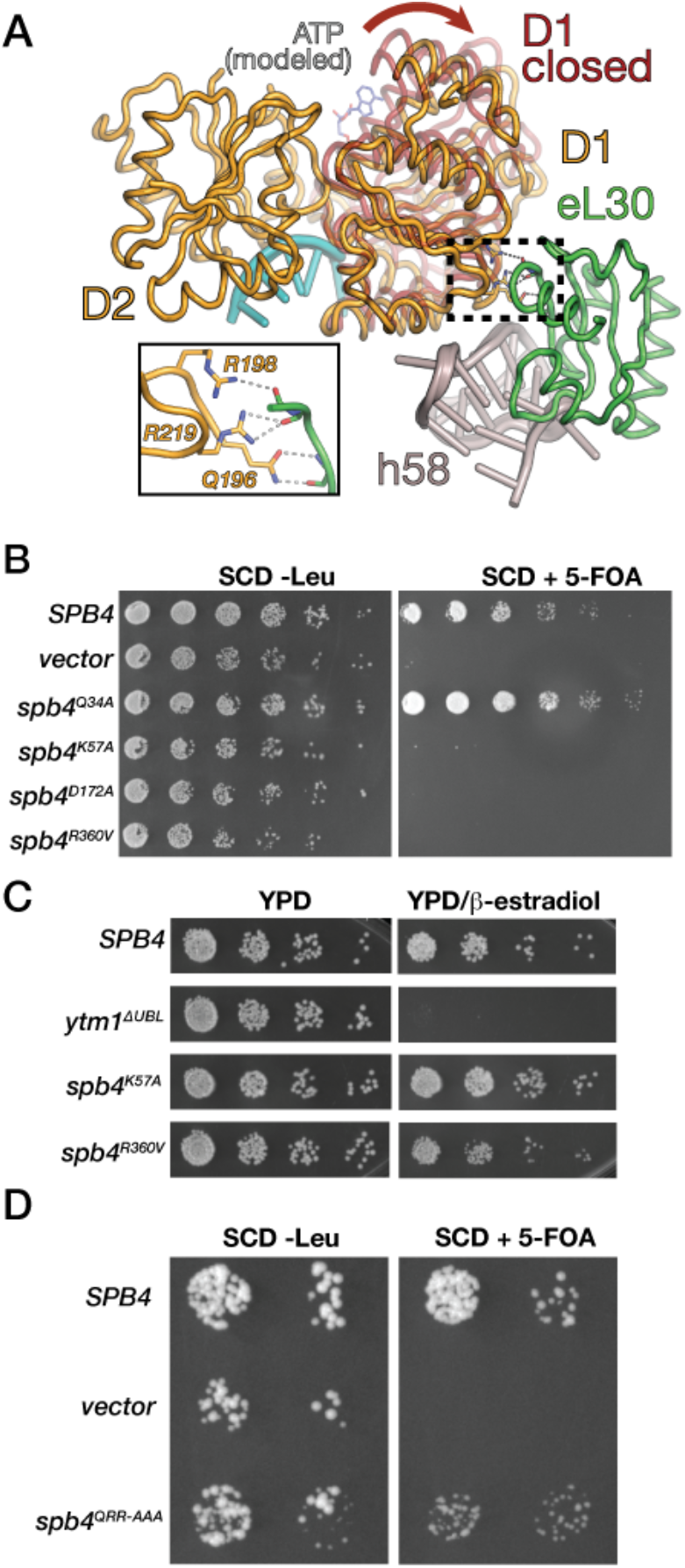
ATP binding and hydrolysis by Spb4 are essential for the formation of the high-energy intermediate. (A) Cartoon representation of the Spb4-eL30-h58 interface, inset shows residues at the interface with hydrogen bonds shown as dashed lines. A hypothetical model of Spb4 D1 in a closed state is superposed on our model and is colored maroon to show the ∼5° swivel motion of D1 relative to D2 observed in the high-energy intermediate. A stick model of ATP is shown based on its binding site in the closed conformation. (B) Plasmid shuffle assays testing the ability of *SPB4* catalytic mutants to rescue the lethality of *spb4Δ*. Left panel – serial dilutions of cells grown at 30° in the presence of pSPB4-URA3 and the LEU2 plasmids listed. Right panel – serial dilutions of mutant strains after counterselection against pSPB4-URA3 on 5-FOA plates. (C) Estradiol induced overexpression of the ATP binding mutant *spb4*^*K57A*^ and the ATP hydrolysis mutant *spb4*^*R360V*^, *ytm1*^*ΔUBL*^ is shown as a positive control. (D) Plasmid shuffle assay showing that disruption of the Spb4/eL30 interface (*spb4*^*QRR-AAA*^) leads to a moderate growth defect on 5-FOA plates (right panel – compare colony sizes).

## The high-energy Spb4 intermediate promotes rRNA domain IV root helix organization and the recruitment of the Spb1-methyltransferase domain

Having established the rRNA remodeling function of Spb4 and defined the structural details of its trapped post-catalytic conformation, we set out to understand the functional role of this high-energy intermediate in 60S ribosome maturation. We therefore focused on the differences between the sequential E1 and E2 states, as well as comparisons to the NE1 state. In states E1 and E2, the overall structure of the h62/h63/ES27 region is similar (Fig. 5, A and B). However, h64 and much of root helix h61 are disordered in state E1 before organizing into a helical stack in state E2. In addition to rRNA organization, the transition between E1 and E2 results in specific RBF binding events. The E1 state shows partial density for an N-terminal domain of Spb1, which we term the anchoring domain (Spb1-AD, residues 254-329). In E2, the MTD of Sbp1 (residues 1-222) binds the cleft between helices h92 and h64, helping stabilize the newly formed h64/61 junction (Fig. 5, B and C). In addition, E2 contains a previously unresolved linker region between Sbp1-AD and Sbp1-MTD (residues 223-253) that engages the h64-h61 junction with two conserved arginines, simultaneously stabilizing the root-helix junction and promoting Spb1-MTD binding (Fig. 5C). Finally, Rrp17 binds opposite to Sbp1-MTD on h64 and forms an elongated helical structure that acts as a structural bracket between Spb1-MTD and Spb4-D1 (Fig. 5, B and D). Spb1-MTD engagement leads to the 5’-O-ribose methylation of nucleotide G2922, and our improved maps allow for a complete atomic model of the Spb1-MTD/rRNA interaction to be built, showing that Spb1-MTD remains bound post-catalytically, as the methyl group on G2922 and the presence of SAH in the Spb1-MTD active site are clearly defined in our maps. These data support a model in which formation of the h61/h64 junction requires formation of h62-alt by Spb4 and is stabilized by the cooperative binding of the Sbp1-MTD to the newly organized domain IV root structure.

**Figure 5.**
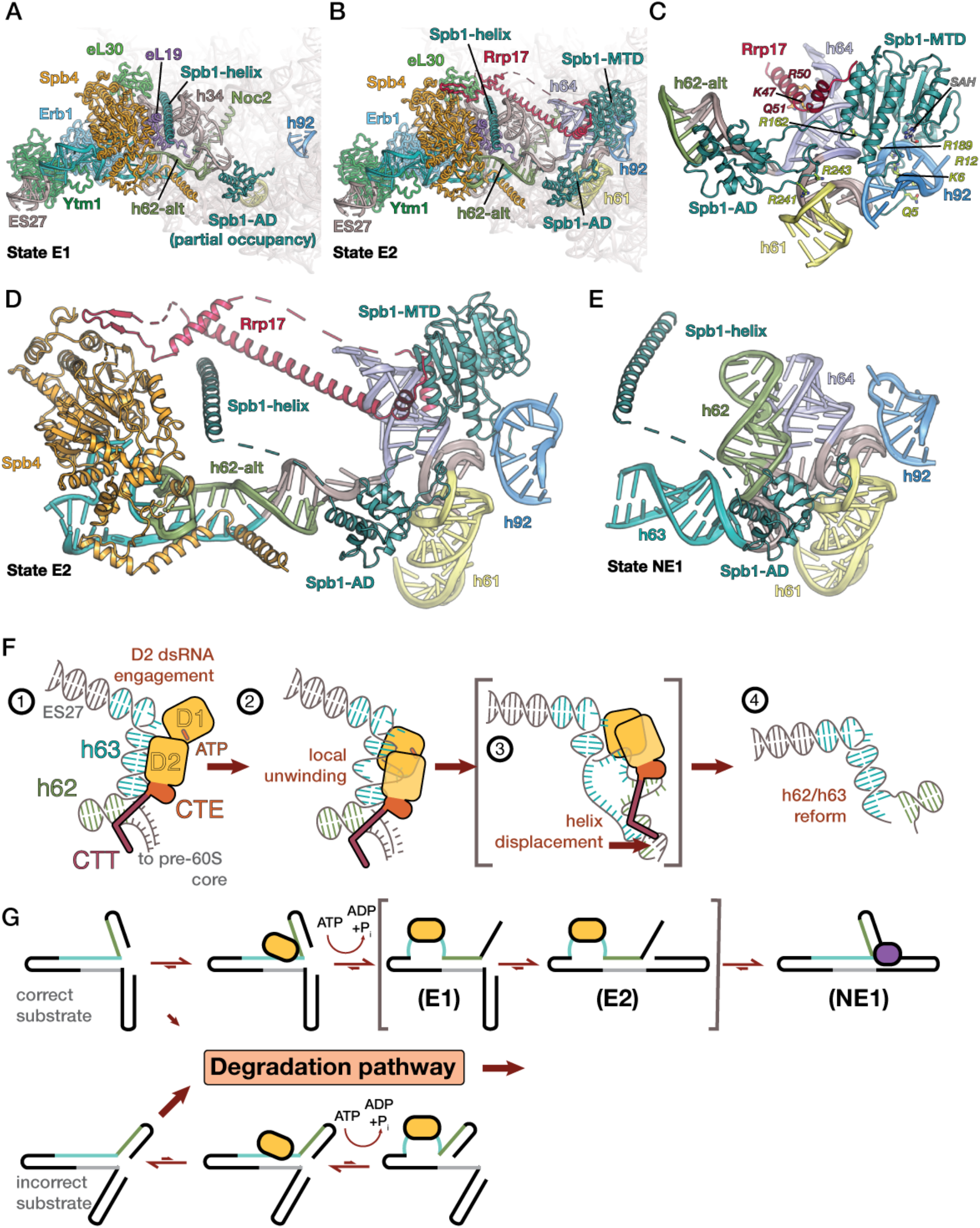
The Spb4 high-energy intermediate drives the assembly of the domain IV rRNA root helix. Cartoon representation of the compositional difference between the sequential E1 (A) and E2 (B) states, showing the organization of the h61 domain IV root helix and subsequent binding of the Spb1-MTD and of Rrp17. (C)Detailed cartoon view how Spb1-MTD binds the domain IV root junction. The linker domain between Spb1-AD and Spb1-MTD directly engages the h61/h64 junction with a pair of ARG residues (R241/R243 – shown as green sticks). Spb1-MTD is wedged between h92 (which contains the G2922 nucleotide substrate) and h64, making specific contacts (shown as green sticks) with its N-terminal tail and internal loop regions. Rrp17 (shown in red) also makes specific contacts (shown as red sticks) with h64. (D) Simplified cartoon representation of state E2, showing how the RBF Rrp17 bridges Spb1 and Spb4 in the trapped high-energy intermediate, connecting Spb4-directed rRNA remodeling with the organization of the domain IV root helix structure. This is the state immediately prior to Spb4 release. (E) Simplified cartoon representation of state NE1 in the same orientation as state E2 in panel D. Spb4 release causes the unwinding of h62-alt and the *in situ* reformation of rRNA helices h62 and h63, completing the mature conformation of the h61 root helix structure. (F) Schematic of the mechanism of sequence-directed rRNA remodeling by Spb4. 1. Spb4 D1 and D2 engage ATP and dsRNA respectively. 2. The D1/D2 interface is formed, leading to the local unwinding of h63 and engagement of ssRNA at the D1/D2 domain interface. 3. The local unwinding in the previous step is expanded through sequence-specific interactions with the Spb4-CTE and made energetically favorable by the formation of h62-alt, forming the high-energy Spb4-containing intermediates described in this study. 4. After removal of Spb4, h62/h63 revert to their most stable conformation, locking in the conformational changes facilitated by the formation of h62-alt. (G) Kinetic proofreading model of 60S nucleolar biogenesis. Schematic assembly path showing Spb4 engagement of correct (top) and incorrect (bottom) RNP substrates (adapted from (*16*)). The sequence and context specificity of Spb4 activity heavily favors high-energy intermediate stabilization (shown in brackets – states E1 and E2) and forward progress (State NE1) only with a correct RNP substrate. DEAD-box ATPase engagement of incorrect RNPs without precisely aligned complimentary strands will self-resolve without forming a high-energy intermediate. These intermediates become kinetically trapped and can be targeted for degradation.

## Spb4 dissociation releases the high-energy intermediate to lock in the mature domain IV root structure

Because initial rRNA engagement and unwinding by Spb4 is too transient and dynamic for structural analysis, we instead focused on the disassembly mechanism of the Spb4 high-energy intermediate for further insights. Recent studies have shown that, immediately after Rea1-catalyzed removal of the Ytm1/Erb1 complex and its associated factors, including Spb4, a transitional state characterized by the presence of both Spb1 and Nop53 can be isolated (*24*). We obtained an improved reconstruction of this state by isolating particles from a strain with dual tags on Spb1 and Nop53 (Fig. 1, C and D, fig. S2B). This reconstruction, with a resolution of 2.8Å, allowed for a complete molecular model of the h62/h63/ES27 region to be built, representing the structure of the pre-60S immediately after Spb4-release and h62-alt disassembly (Fig S3, L to N and fig S6.). We observe that, in addition to binding the RBF Nop7 in a manner analogous to Erb1, Nop53 possesses an internal helix that helps stabilize ES27 in its near-mature conformation (Fig. 2D and fig. S5F) prior to the recruitment of the RBF Arx1. While we do not see strong density for the Spb1-MTD, Spb1-AD is still present, as is the central helix of Spb1 (Fig. 5E). The retention of these structures means that the spontaneous reformation of h62 and h63 occurs in a tightly constrained space upon the release of Spb4, consistent with our interpretation that the interaction network surrounding Spb4 stabilizes a high-energy rRNA structural intermediate that snaps back into its most stable conformation after Spb4 release. Importantly, although h62 and h63 reform, they do so in a manner that preserves the mature configuration of the domain IV root structure (Fig. 5, D and E). The organization of domain IV now allows for the incorporation of the L1 stalk structure into the core of the 60S, providing a clear mechanism that imposes directionality to this step in 60S biogenesis (*24*).

## Sequence-directed helix remodeling expands the functional repertoire of DEAD-box ATPases

Our structural and genetic studies reveal that the DEAD-box ATPase Spb4 triggers the sequence-directed remodeling of an internal, topologically restrained RNA structure by leveraging a small, localized strand unwinding event into a broad RNA restructuring within the maturing RNP (Fig. 5F). The thermodynamic cost of strand opening within the constrained substrate is offset by the presence of a position- and sequence-specific RNA element flanking the primary dsRNA engagement site. The presence of this RNA element allows Spb4 to stabilize an alternate, thermodynamically less favored base-pairing arrangement to yield a high-energy pre-60S intermediate (Fig. 5F). This mechanism explains both how the limited strand separation activity of DEAD-box ATPases can be harnessed for restructuring RNPs and how substrate specificity is achieved by structural elements flanking the catalytic core of these proteins. Indeed, a subset of DEAD-box ATPases involved in ribosome biogenesis contain CTEs that are structurally similar to Sbp4, suggesting that CTEs in these enzymes also define their sequence specificities (Fig. S7A). Consistent with this, residues responsible for direct base interactions in Spb4 are not conserved among these DEAD-box ATPases (Fig. S7B).

In yeast, 14 DEAD-box proteins are required for 40S and 60S ribosome biogenesis. We propose that a subset of these enzymes may also act as sequence-guided RNA remodeling factors in a manner similar to Spb4. The wider prevalence of such activities predicts the existence of complementary sequences near the initial strand separation sites of other DEAD-box ATPases. While the binding sites and substrate specificities of the other DEAD-box ATPases involved in ribosome maturation remain unknown, alternative base-pairing arrangements have been proposed, most notably between ITS-1 and the 5’ end of the 5.8S rRNA(*17*)}. The mechanism exemplified by Spb4 provides a mechanistic framework to identify and characterize other cryptic rRNA complementary sequences in ribosome biogenesis and other RNP remodeling events modulated by DEAD-box ATPases.

While all previous structures of RNA-bound DEAD-box proteins rely on nucleotide for stability, Spb4 remains bound to its product after ATP hydrolysis and ADP release, instead relying on *trans* interactions to maintain the high-energy RNA intermediate (Fig. 3E, fig. 4A). Our structures do not reveal how ATP hydrolysis is triggered, nor are the advantages of trapping Spb4 in the apo state obvious. We hypothesize that early nucleotide hydrolysis and release ensures that Ytm1, Erb1 and Spb4 disengage the unwound portion of h63 simultaneously to rapidly trigger the unwinding of h62-alt and reformation of h63 and h62, optimizing the docking kinetics of the mature form of h62 onto the pre-60S core (Fig. 5, D, E and F).

## DEAD-box ATPase function and kinetic proofreading of ribosome biogenesis

Internal quality control mechanisms have long been postulated for ribosome biogenesis and DEAD-box ATPases named as candidates (*17, 36*). We believe that Spb4-triggered RNA remodeling exemplifies a high-energy intermediate state as predicted by this model (Fig. 5G). This intermediate is maintained until additional RPs and RBFs bind and stabilize the domain IV root helix, “locking in” its final rRNA tertiary structure and ensuring directional progress (Fig. 5). In contrast, misassembled substrates cannot be rapidly and irreversibly remodeled, therefore becoming kinetically trapped and targeted for degradation (Fig. 5G) (*16*). Spb4 is the last of a cluster of DEAD-box ATPases that engage the nucleolar pre-60S in rapid succession (*22*). The passage of assembly intermediates through sequential surveillance checkpoints defined by individual rRNA remodeling events could act as a “molecular ratchet” (*16*) to increase the overall accuracy of the 60S maturation process.

## Acknowledgements

We thank Michael Rosen, Luke Rice, James Berger and Xiaochen Bai for helpful discussions and Matthew Parker for help with live-cell imaging. We thank Daniel Stoddard at the UTSW Cryo-Electron Microscopy Facility, James Chen at the UTSW Structural Biology Lab and Theo Humphreys at the Pacific Northwest Center for Cryo-EM.

The UTSW Cryo-Electron Microscopy Facility is funded in part by the CPRIT Core Facility Support Award RP170644.

A portion of this research was supported by NIH grant U24GM129547 and performed at the PNCC at OHSU and accessed through EMSL (grid.436923.9), a DOE Office of Science User Facility sponsored by the Office of Biological and Environmental Research.

This work was supported by funding from:

The German Research Foundation (STE 2517/1 and STE 2517/5-1) (FS)

The Collaborative Research Centre 969 (Project A06) (FS)

The Cancer Prevention and Research Institute of Texas (RR150074) (JPE),

The Welch Foundation (I-1897) (JPE)

The UTSW Endowed Scholars Fund (JPE)

The National Institutes of Health (GM135617-01) (JPE)

## Author contributions

Conceptualization: VEC, CSW, JPE

Methodology: VEC, CSW, NP, SB, JPE

Resources: CS, CSW, SB, VEC

Investigation: VEC, KS, NP, CSW, SB, CS

Writing - original draft: JPE and VEC.

Writing - review & editing: JPE, VEC, CSW, KS, CS, FS

Visualization: VEC, JPE

Supervision: FS, JPE

Funding Acquisition: FS, JPE

## Competing interests

The authors declare no competing interests.

## Data and materials availability

The cryo-EM density maps and models have been deposited in EMDB and PDB with accession codes: EMD-24296/pdb-7R7A (E1 Overall map/model), EMD-24290/pdb-7R72 (E1 Spb4 local map/model), EMD-24269/pdb-7NAC (E2 Overall map/model), EMD-24270/pdb-7NAD (E2 Spb4 local map/model), EMD-24271/pdb-7NAF (E2 Spb1 local map/model), EMD-24280/pdb-7R6K (E2 Noc2/Noc3 local map/model), EMD-24297/pdb-7R7C (E2 L1 stalk local map/model), EMD-24286/pdb-7R6Q (E2 foot local map/model), EMD-26259/pdb-7U0H (NE1 Overall map/model). Yeast strains, plasmids and other materials are available from J.P.E. upon request.

## Supplementary Materials

Materials and Methods

Figs. S1 - S7

Tables S1 - S3

## Notes

### Competing Interest Statement

The authors have declared no competing interest.

