## Supplemental information for "Sequence-directed RNA remodeling within a topologically complex RNP substrate"

### SUPPLEMENTAL MATERIALS

#### Materials and Methods

Yeast Strains and Media

All yeast strains used in this study are listed in table S1, all plasmids used in this study are listed in table S2. Genomic knockout strains were generated as described previously (*37*) (*38*).

Microscopy

Strains were grown overnight in YPD to saturation, then diluted 20-fold into SC-Trp with β-estradiol to a final concentration of m for 1.5 hours. Images were taken on a Nikon Ti2-E inverted microscope equipped with a Yokogawa CSU-X1 spinning disk and a 100x NA = 1.49 oil objective. Samples were illuminated with a 488 nm solid state laser light source and images collected on an ORCA-FLASH 4.0 sCMOS camera. A z-series of confocal images was acquired for two fields of view in a single biological replicate. Image processing was performed using ImageJ.

Overexpression and Purification of pre-60S Particles

Starter cultures were grown overnight in YPD to saturation. The following morning, 12L of YPD were inoculated from the starters to an OD600 of 0.1 and grown at 30ºC to an OD600 of 0.8. For overexpression of tagged Ytm1ΔUBL and Spb4, cells were induced for 1 hour by the addition of β-estradiol to a final concentration of 2 µM. Cells were harvested by centrifugation for 20 min at 4000x g. Cell pellets were washed in Ribo-buffer A (50 mM Bis-Tris KCl pH 8.0, 150 mM NaCl, 10 mM MgCl2, 1 mM TCEP, 0.1% (w/v) NP-40) and centrifuged for 10 min at 4000 g, then harvested and flash frozen. Cryogenic lysis was done using a grinding ball mill (Fritsch Pulverisette 6). For purification of Ytm1ΔUBL·Spb4 pre-ribosomes, 20 g of lysate was thawed and resuspended in Ribo-buffer A supplemented with E64, pepstatin and PMSF. Lysate was cleared by centrifuging at 100,000 g for 30 min and loaded onto IgG Sepharose resin (Cytiva). Samples were washed with Ribo-buffer A, followed by Ribo-buffer B (50 mM Bis-Tris KCl pH 8.0, 150 mM NaCl, 10 mM MgCl2, 1 mM TCEP and 0.01% (w/v) NP-40). Cleavage of protein-A was done on-column using 3C protease, eluates were applied to a Strep-Tactin column (Cytiva) and washed with Ribo-buffer B. Elution was done by on-column cleavage of Strep using bdNEDD8 protease. For purification of Nop53·Spb1 pre-ribosomes, the sample was obtained from steady-state cells grown in YPD to an OD600 of 0.8. Cell harvesting, lysis and purification was performed as described above, except that the elution from the Strep-tactin column was performed by flowing 5 column volumes of Ribo-buffer B enriched with 10 μM desthiobiotin over the column. In both cases eluates were concentrated using Amicon ultra 0.5 ml spin columns with 100 kDa cutoff (Merck Millipore) for cryo-EM analysis.

Grid Preparation and Data Collection

Cryo-EM grids were prepared by applying 3 $\mu$L of Ytm1ΔUBL·Spb4 samples at a concentration of 4.5 A260ml-1 or Spb1·Nop53 at 2.0 A260ml-1 to glow-discharged continuous carbon coated Quantifoil R2/2 300-mesh or Quantifoil R2/1 300-mesh grids respectively. Grids were blotted for 3.5 s with a blot force of 12 and wait time of 15 s under 100% humidity at 4 ºC followed by plunge freezing into liquid ethane using a Mark IV Vitrobot (FEI).

Micrographs were acquired on a Titan Krios (FEI) operated at 300 kV equipped with a Gatan K3 direct electron detector using a slit width of 30 eV on a GIF-Quantum energy filter. Automated data collection was performed using SerialEM (*39*) with a defocus range of 0.9-2.2 $\mu$m. For Ytm1ΔUBL·Spb4 grids, each micrograph was dose fractionated into 50 frames of 0.05 s each under a dose rate of 26.2 e-/pixel/s with a pixel size of 1.08 Å, a total exposure time of 2.5 s and a total dosage of about 65.5 e-/pixel. For Spb1·Nop53 grids, each micrograph was dose fractionated into 40 frames of 0.05 s each under a dose rate of 26.2 e-/pixel/s with a pixel size of 1.02 Å, with a total exposure time of 2 s and a total dosage of about 52.4 e-/pixel.

Cryo-EM data processing

Motion correction was performed using MotionCorr2(*40*), and CTF parameters were estimated using GCTF(*41*). All subsequent image processing was performed using RELION (*42*). Initially, about 300 particles from 3 micrographs were extracted and binned to generate initial 2D classes as templates for automated particle picking. Automatically picked particles were extracted and binned 4 times for 2D classification. 2D classes resembling pre-ribosomes and showing clear secondary structure features were selected for 3D classification. Reference maps for 3D classification were low pass filtered to 15 Å. 3D classes were inspected in Chimera, and selected 3D classes were combined and re-extracted at the original pixel size of 1.08 Å. 3D refinement was performed with an imposed symmetry of C1, followed by CTF refinement and another cycle of 3D refinement with postprocessing. For localized classification and refinement a mask with a soft edge was created around the site of interest, the core was subtracted and followed by a round of local 3D classification *(43)*. Particles that showed strong occupancy for the desired region were selected for 3D refinement. For state E2, overlapping local masks centered on the Spb1-MTD (3.13 Å), Spb4 (3.04 Å), Noc2/Noc3 (3.17 Å), the "foot" (2.98 Å) and the L1 stalk (3.71 Å) were used to obtain locally refined maps. An overall (3.04 Å) and locally refined map with a mask around Spb4 (3.07 Å) was also obtained for the particle subset lacking Spb1-MTD (State E1). Local maps generated for the Spb1·Nop53 particle (State NE1) were used to select a homogeneous subset of particles for final refinement, the local maps were not used for model building. The final resolution in each case was estimated by applying a soft mask over the maps and was calculated using the gold-standard Fourier shell correlation (FSC) = 0.143(*44*). Local resolution maps were obtained using ResMap (*45*).

<u>Model Building and Refinement</u>

PDB 6ELZ was used as a starting model(*25*) for states E1 and E2. Spb4 was built *de novo* starting from a Phyre2 *(46)* model that contained the core RecA domains, the CTE and CTT of Spb4 were built manually into the density. PDB 6YLX was used as a starting model for state NE1 (*24*). Interpretation of density was facilitated and validated with XL-MS data (*22*) (*47*). All models were built in Coot (*48*) and refined in PHENIX using phenix.real_space_refine (Table S3) (*49*). Figures were generated with UCSF Chimera (*50*), Chimera X *(51)*, and PyMOL Molecular Graphics System (Version 2.0 Schrödinger LLC.).

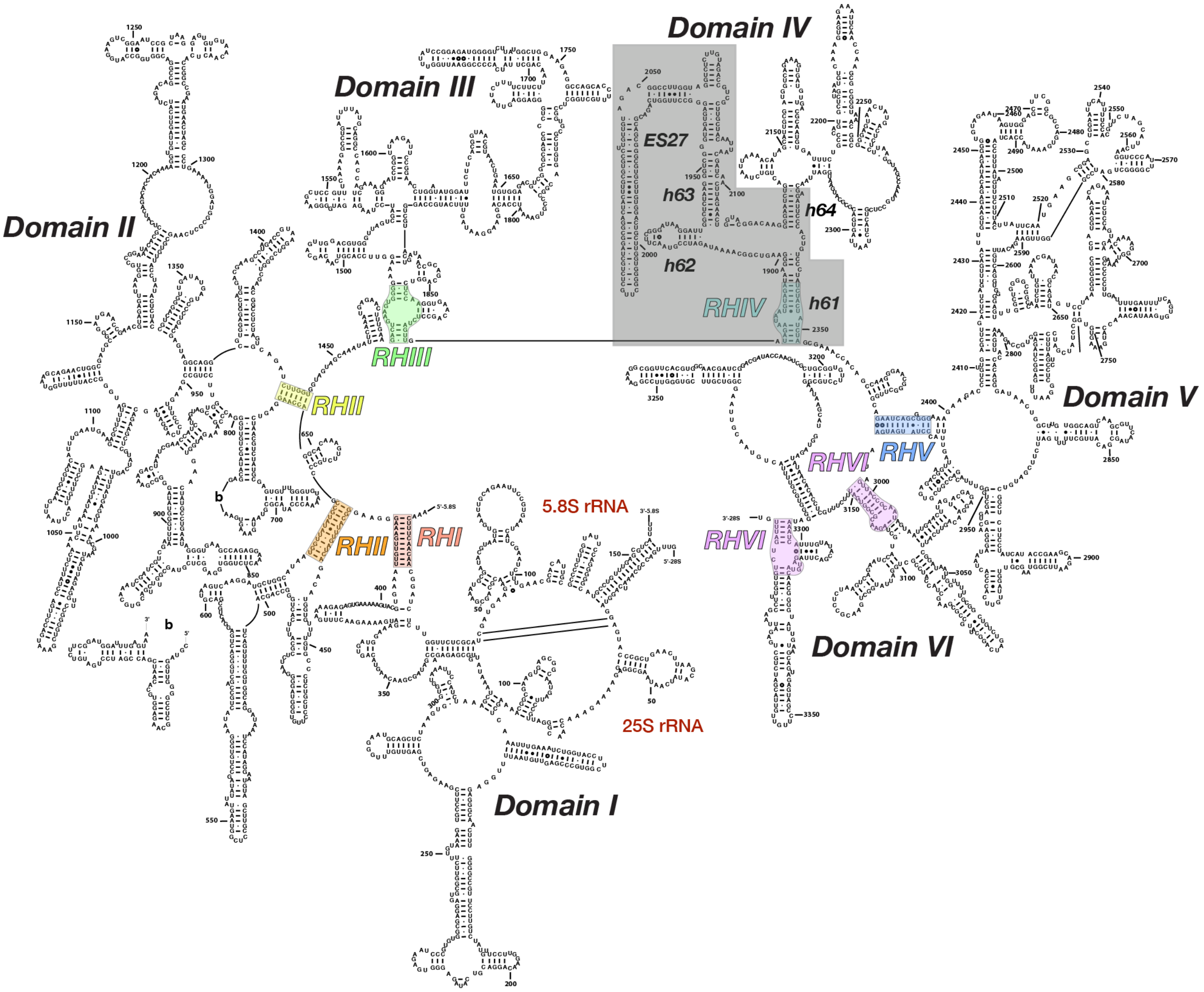


**Figure S1. Position of the domain IV elements engaged by Spb4 within the secondary structure of the mature 25S rRNA.**

Secondary structure diagram of the mature 25S and 5.8S rRNA, showing the major subdomains with their root helices highlighted and labeled. The region in domain IV that is remodeled by Spb4 is boxed and important rRNA helices labeled.

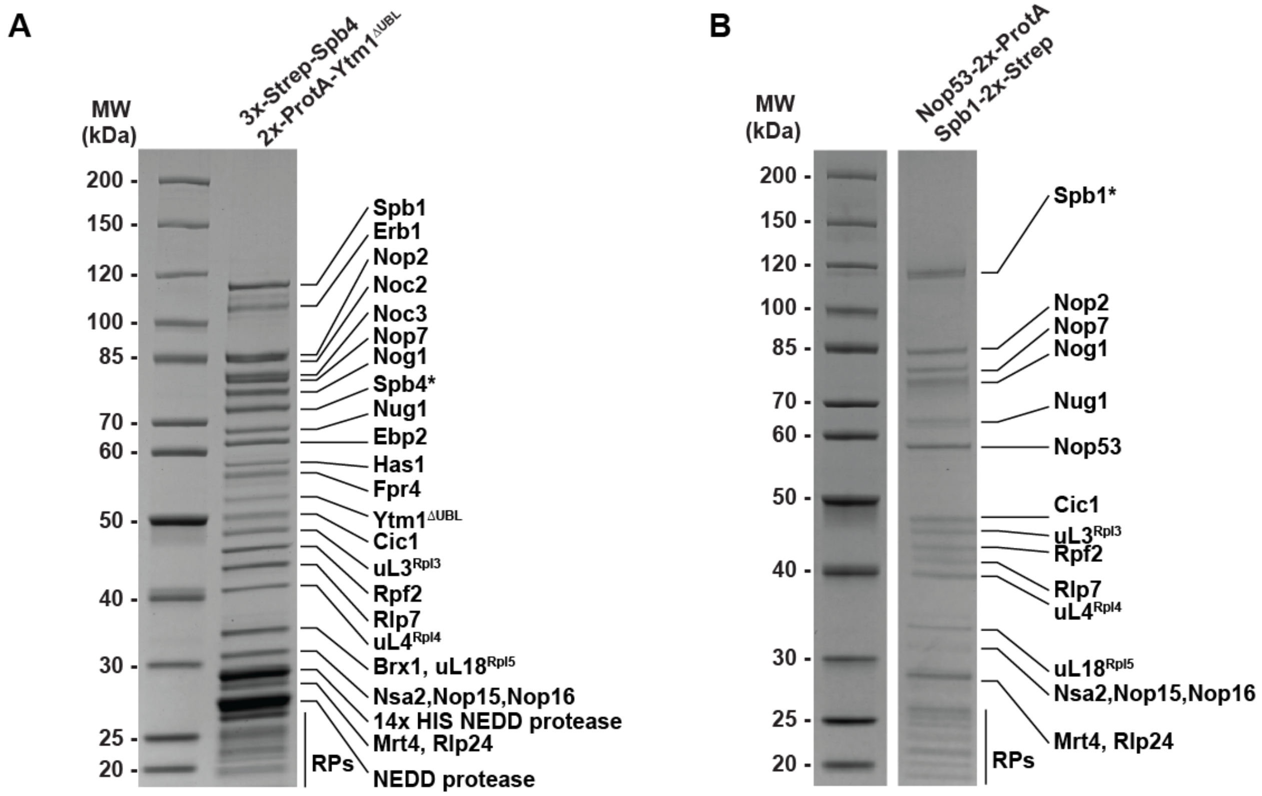


**Figure S2. Purified samples of Ytm1$^{ΔUBL}$/Spb4 and of Nop53/Spb1 particles used for Cryo-EM single particle reconstruction.**

4-20% SDS-PAGE Gel of purified (A) Ytm1$^{ΔUBL}$/Spb4$^{WT}$ or (B) Nop53/Spb1 pre-ribosomal intermediates. Protein identification was determined based on molecular weight, ambiguous bands were identified by mass spectrometry.

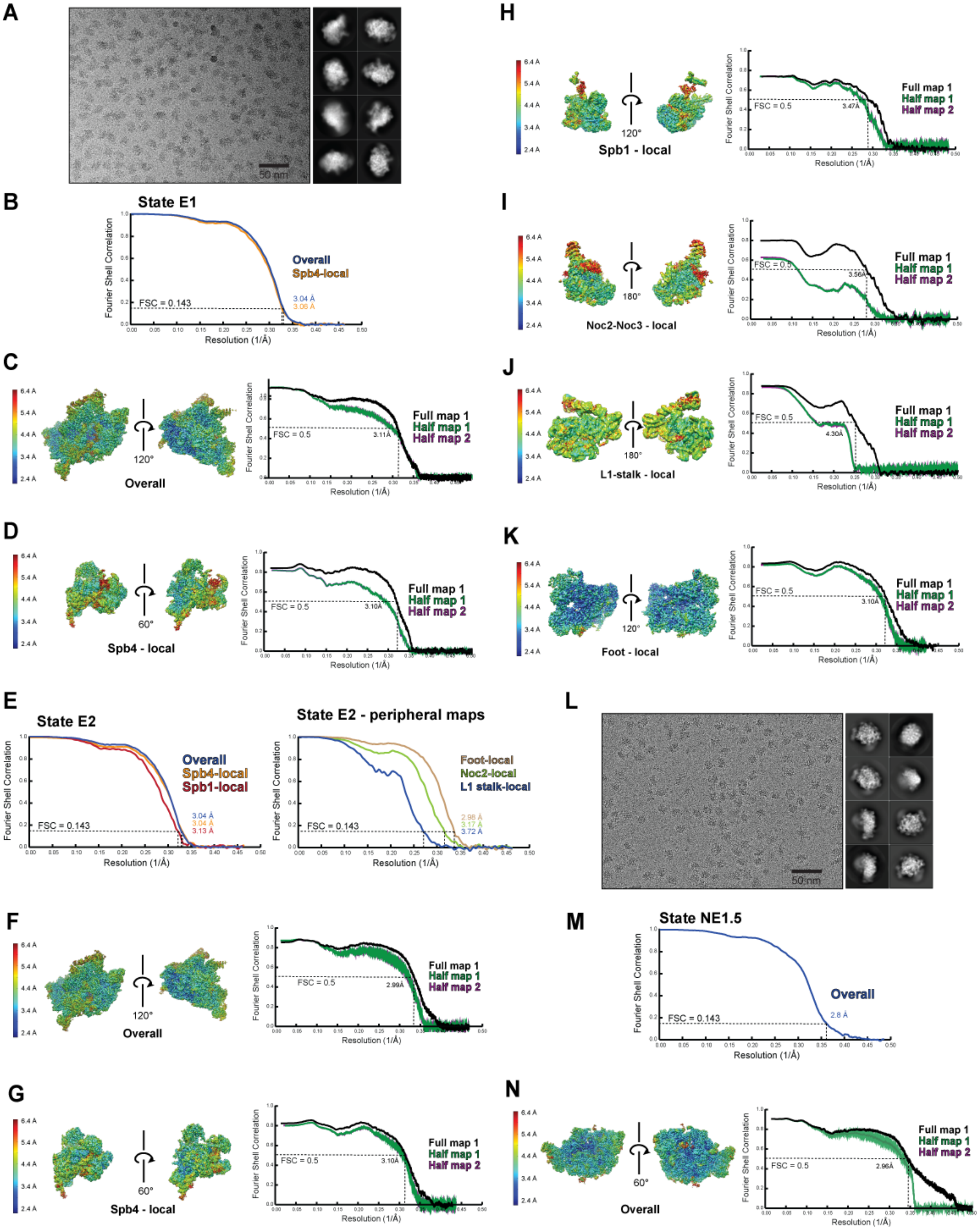


**Figure S3. Micrographs, 2D classes and Fourier shell correlation (FSC) curves for reconstructions and model validation.**

(A) Representative electron micrograph and 2D classes of the Ytm1$^{\Delta UBL}$/Spb4$^{WT}$ sample that yielded States E1 and E2.

(B) FSC curves for the overall and Spb4-focused maps corresponding to State E1.
(C – D) Local resolution filtered maps (left) and model to map FSC curves for each half map and full map (right) for the overall and Spb4-focused local map for State E1.
(E) FSC curves for the Overall, Spb4-focused and Spb1-focused map from State E2 (left). FSC curves for the peripheral maps focused on the foot, Noc2, and the L1-stalk from State E2 (right).
(F – K) Local resolution filtered maps (left) and model to map FSC curves for each half map and full map (right) for the overall and local map corresponding to State E2.
(L) Representative electron micrograph and 2D classes of the Spb1/Nop53 sample that yielded State NE1.
(M) FSC curves for the Overall map corresponding to State NE1.
(N) Local resolution filtered map (left) and model to map FSC curves for each half map and full map (right) for the overall map corresponding to State NE1.

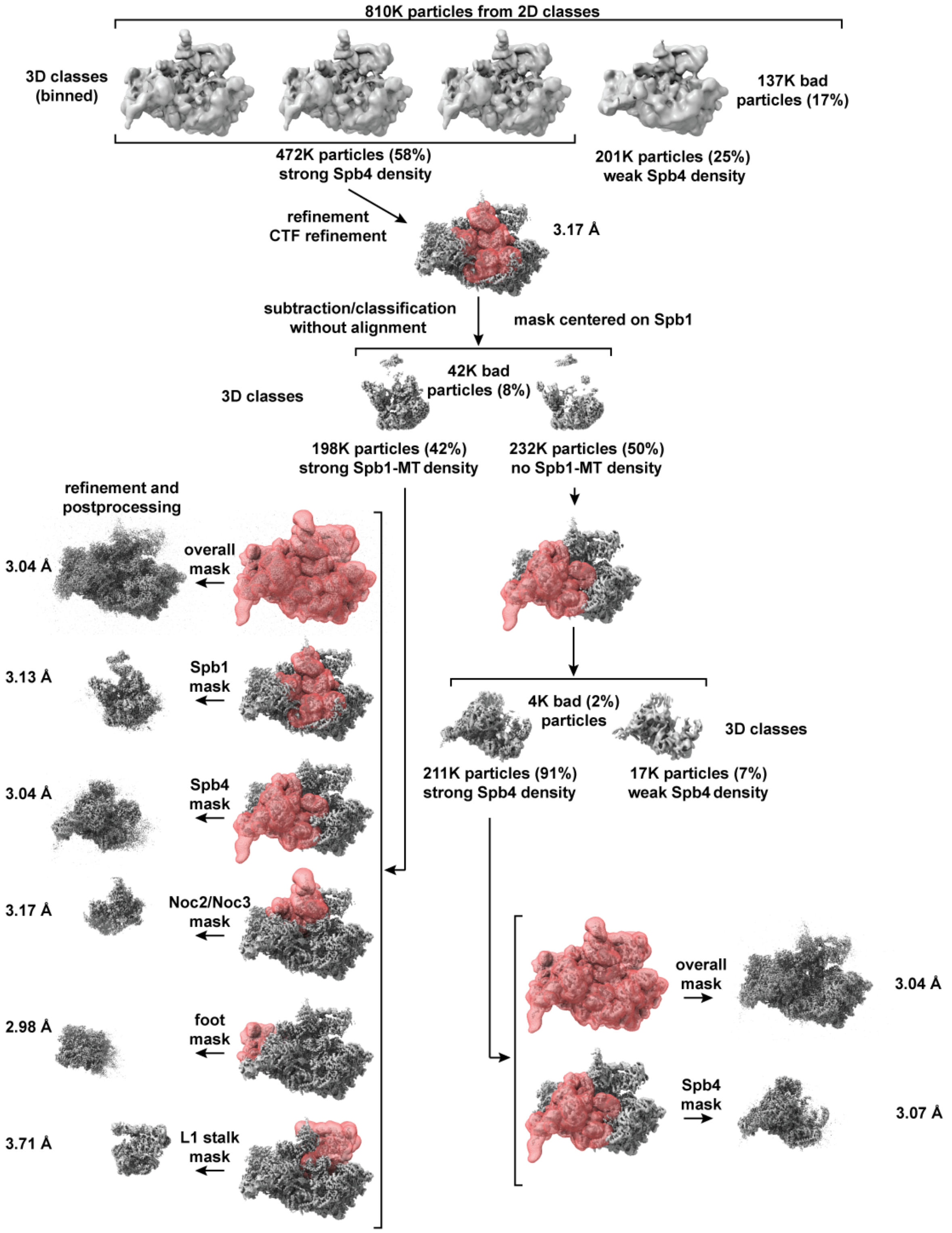


**Figure S4. Data processing, classification and sorting scheme for State E1 and State E2.** Work flow of the data processing strategy after 2D classification. Map volumes and masks (red) are shown for all critical steps. Major sorting and classification criteria, particle numbers and percentages (for each 3D classification) as well as final map resolutions are indicated.

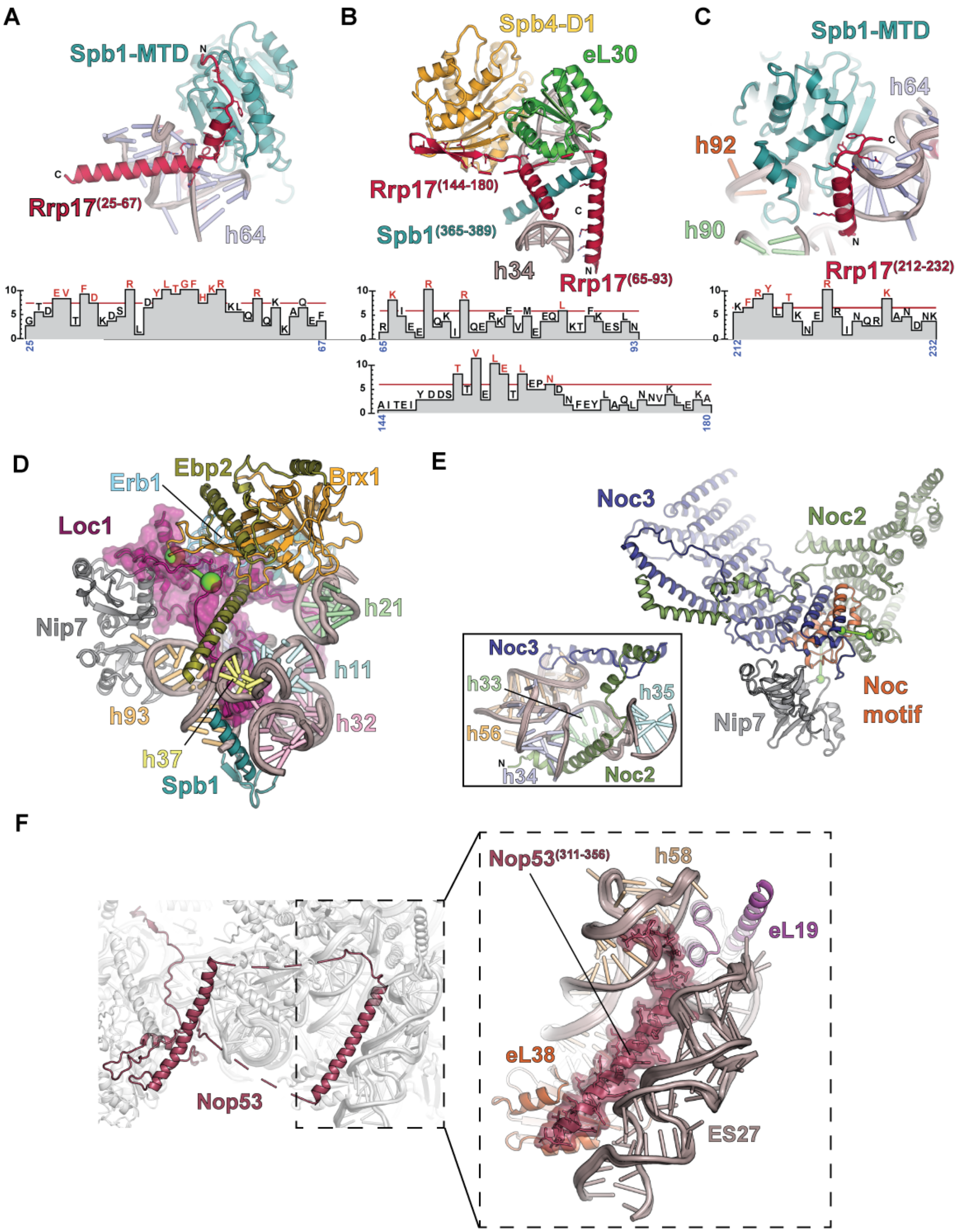


**Figure S5. Interactions of the biogenesis factors Loc1, Noc2 and Rrp17 in state E2, and of the newly modeled helix of Nop53 in state NE1.**

(A) Cartoon representation of the interaction between Rrp17 (red), Spb1-MTD (teal) and h64 (blue). A plot of residue conservation scores based on an alignment of 15 fungal Rrp17 homologs (*52*) is shown below the model. Residues with a conservation score >7 are highlighted and shown as sticks in the structural model.

(B) Cartoon representation of the interaction between Rrp17 (red), eL30 (green), Spb4-D1 (yellow), the Spb1 helical element (teal) and h34 (gray). Conserved residues (identified in plot) are shown as sticks (same criteria as in panel A).
(C) Cartoon representation of the interaction between the C-terminal helix of Rrp17 (red), Spb1-MTD (teal) and h64 (blue). Adjacent rRNA helices h90 (green) and h92 (orange) are also shown. Conserved residues (identified in plot) are shown as sticks (same criteria as in panel A).
(D) Detail of the position and interactions of Loc1 within the state E2 60S particle. The cartoon representation shows the interaction of Loc1 (purple) with various RBFs and rRNA elements. Model building and validation was aided by a crosslink between Loc1 and Brx1 (green connected spheres) (*22*).
(E) Cartoon representation of the Noc2 (green)/Noc3(blue) interaction highlighting the conserved Noc module (orange) of Noc3 and the extended N-terminus of Noc2, which extends into the 60S core (inset). Positions of crosslinks between Noc2 and Noc3 and Noc2 and Nip7 are shown as green connected spheres (*22*). The interaction between Noc2/Noc3 is analogous to the interaction between Noc4 and Nop14, revealing Noc2 and Nop14 to be structural paralogs(*53*).
(F) State NE1 with Nop53 colored maroon for emphasis. The newly modeled Nop53 helix is boxed and expanded in the inset. Surface view of the helix is shown for clarity.

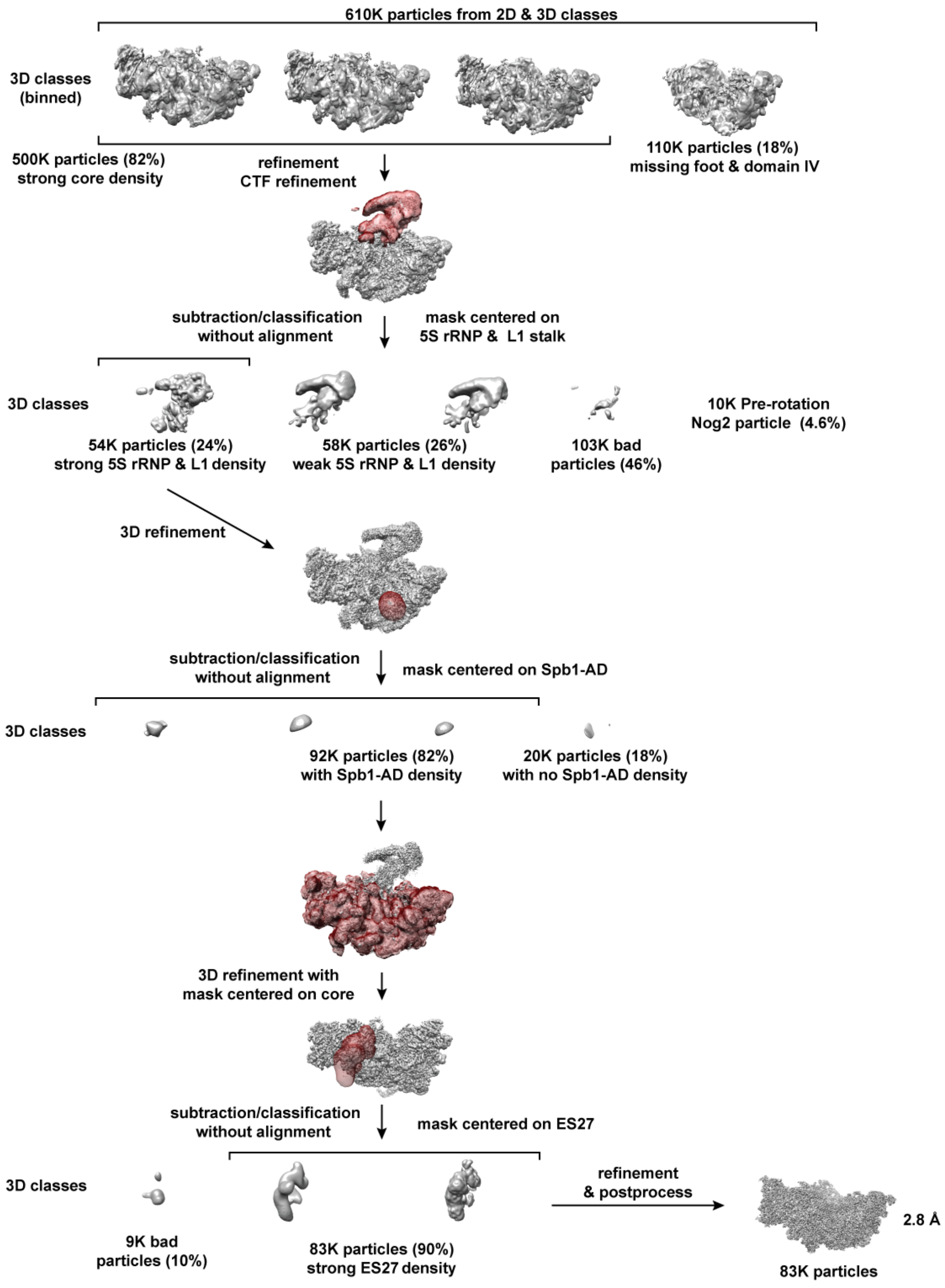


**Figure S6. Data processing, classification and sorting scheme for State NE1.**
Work flow of the data processing strategy after 2D classification. Map volumes and masks (red) are shown for all critical steps. Major sorting and classification criteria, particle numbers and percentages (for each 3D classification) as well as final map resolutions are indicated.

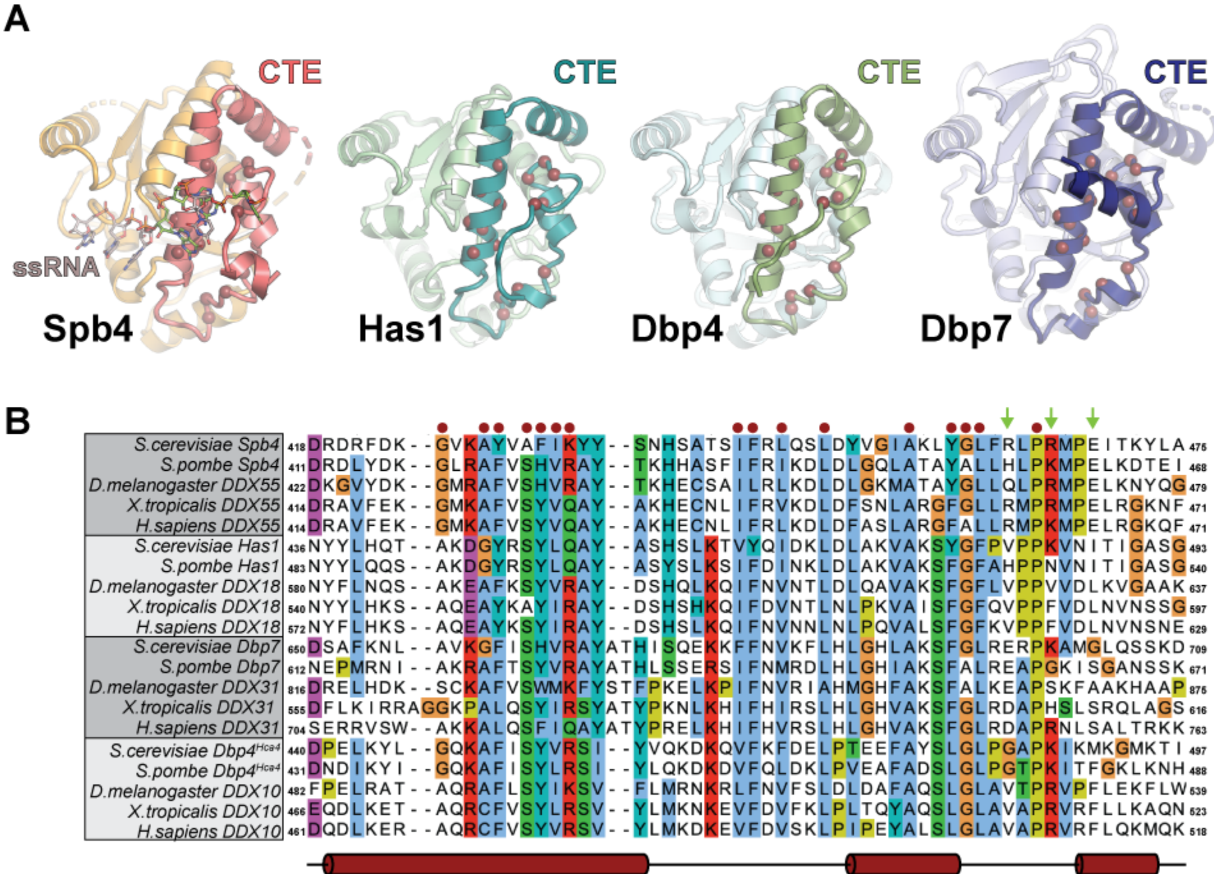


**Figure S7. Structural homology to the Spb4-CTE defines a subset of DEAD-box ATPases involved in ribosome biogenesis.**

(A) Comparison of the CTE of Spb4, Has1 (pdb 6C0F (*54*)), Dbp4 (Alphafold model) and Dbp7 (Alphafold model) (*55*). Red spheres represent conserved residues among DEAD-box ATPases (B) Multiple sequence alignment of the CTE of Spb4, Has1, Dbp7 and Dbp4. The helix boundaries of Spb4 and Has1 are shown as red tubes and conserved residues, also shown in the structure cartoons, are marked with red dots. Residues implicated in base specific interactions are not conserved and indicated by green arrows

**Table S1. Yeast strains used in this study**

| Strain | Relevant Genotype | Source |
|---|---|---|
| BY4741 | *MATa his3Δ1 leu2Δ0 met15Δ0 ura3Δ0* | *(56)* |
| YJE201 | *trp1Δ::pACT1-LexA-ER-haB112::TRP1; ura3Δ0::* $P_{LexA\text{-}minCYC1}$*-sfGFP-3xStrep-bdNEDD8-MYC-SPB4::URA3* | this study |
| YJE430 | *trp1Δ::pACT1-LexA-ER-haB112::TRP1; leu2Δ0::* $P_{minCYC1}$*-2xProtA-3C-3xFLAG-ytm1ΔUBL::LEU2; ura3Δ0::* $P_{LexA\text{-}minCYC1}$*-sfGFP-3xStrep-bdNEDD8-MYC-SPB4::URA3* | this study |
| YJE480 | *spb4Δ::HygMX; pRS316-SPB4* | this study |
| YJE599 | *trp1Δ::pACT1-LexA-ER-haB112::TRP1; ura3Δ0* $P_{minCYC1}$*-sfGFP-3xStrep-bdNEDD8-MYC-Spb4(1-510)::URA3* | this study |
| YJE750 | *trp1Δ::pACT1-LexA-ER-haB112::TRP1; SPB1-2xStrep::NatMX; NOP53-3xFLAG-3C-2xProtA::HYG* | this study |
| YJE922 | *trp1Δ::pACT1-LexA-ER-haB112::TRP1; ura3Δ0* $P_{LexA\text{-}minCYC1}$*-sfGFP-3xStrep-bdNEDD8-MYC-Spb4(1-510+NoLS)::URA3* | this study |

**Table S2. Plasmids used in this study**

| plasmid name | description | source |
|---|---|---|
| pAG32 | Template for gene deletion (HygMX) | Addgene |
| pRS315 | Yeast centromeric vector (LEU2) | ATCC |
| pJE718 | pRS315 – SPB4 | This paper |
| pJE731 | pRS315 – SPB4 Q34A | This paper |
| pJE732 | pRS315 – SPB4 K57A | This paper |
| pJE1021 | pRS315 – SPB4 D172A | This paper |
| pJE737 | pRS315 – SPB4 R360V | This paper |
| pJE853 | pRS315 – SPB4 R463A R466A E469A | This paper |
| pJE1042 | pRS315 – SPB4 Δ564 | This paper |
| pJE851 | pRS315 – SPB4 Δ511 | This paper |
| pJE1074 | pRS315 – SPB4 Δ511+ NoLS | This paper |
| pJE1083 | pRS315 – SPB4 Q196A R198A R219A | This paper |
| pRS316 | Yeast centromeric vector (URA3) | ATCC |
| pJE717 | pRS316 – SPB4 | This paper |
| pRS406 | Yeast integration vector (URA3) | ATCC |
| pJE660 | pRS406-$P_{LEXA\text{-}\mathit{minCYC1}}$-sfGFP-3xStrep-bdNEDD8-MYC-*SPB4*-$T_{\mathit{CYC1}}$ | This paper |
| pJE883 | pRS406- $P_{LEXA\text{-}\mathit{minCYC1}}$-sfGFP-3xStrep-bdNEDD8-MYC-*SPB4(1-510)*-$T_{\mathit{CYC1}}$ | This paper |
| pJE1062 | pRS406-$P_{LexA\text{-}minCYC1}$-sfGFP-3xStrep-bdNEDD8-MYC-*SPB4(1-510 + NoLS*)- $T_{cyc1}$ | This paper |
| pNH605 | Yeast integration vector (LEU2) | (*57*) |
| pJE932 | pNH605-$P_{LexA\text{-}minCYC1}$-2xProteinA-3C-3xFLAG-*YTM1*-$T_{ADH}$ | This paper |
| pJE693 | pNH605-$P_{LexA\text{-}minCYC1}$-2xProteinA-3C-3xFLAG-*ytm1ΔUBL* -$T_{ADH1}$ | This paper |

**Table S3. Data collection and refinement statistics**

| | State E1 | | State E2 | | | | | | State NE1 |
|---|---|---|---|---|---|---|---|---|---|
| | Overall | Spb4 local | Overall | Spb1 local | Spb4 local | Noc2/Noc3 local | Foot local | L1-stalk local | Overall |
| **Data collection** | | | | | | | | | |
| Magnification | 81,000x | 81,000x | 81,000x | 81,000x | 81,000x | 81,000x | 81,000x | 81,000x | 81,000x |
| Voltage (kV) | 300 | 300 | 300 | 300 | 300 | 300 | 300 | 300 | 300 |
| Electron exposure ($e^-/A^2$) | 65.5 | 65.5 | 65.5 | 65.5 | 65.5 | 65.5 | 65.5 | 65.5 | 50 |
| Defocus range (mm) | -0.9/-2.2 | -0.9/-2.2 | -0.9/-2.2 | -0.9/-2.2 | -0.9/-2.2 | -0.9/-2.2 | -0.9/-2.2 | -0.9/-2.2 | -0.9/-2.2 |
| Pixel size (Å) | 1.08 | 1.08 | 1.08 | 1.08 | 1.08 | 1.08 | 1.08 | 1.08 | 1.02 |
| Symmetry imposed | $C_1$ | $C_1$ | $C_1$ | $C_1$ | $C_1$ | $C_1$ | $C_1$ | $C_1$ | $C_1$ |
| No. of initial particles | 825,096 | 825,096 | 825,096 | 825,096 | 825,096 | 825,096 | 825,096 | 825,096 | 776,404 |
| No. of final particles | 211,000 | 211,000 | 198,825 | 198,825 | 198,825 | 198,825 | 198,825 | 198,825 | 82,670 |
| Map resolution (Å) | 3.04 | 3.07 | 3.04 | 3.13 | 3.04 | 3.17 | 2.98 | 3.71 | 2.8 |
| FSC threshold | 0.143 | 0.143 | 0.143 | 0.143 | 0.143 | 0.143 | 0.143 | 0.143 | 0.143 |
| **Refinement** | | | | | | | | | |
| Initial model (PDB ID) | 6ELZ | | 6ELZ | | | | | | 3JCT/6YLX |
| Map sharpening B factor (Å$^2$) | -86.35 | -80.3 | -91.67 | -82.77 | -97.16 | -95.57 | -78.23 | -143.14 | -74.53 |
| **Model composition** | | | | | | | | | |
| Non-hydrogen atoms | 153419 | 39994 | 158612 | 35228 | 41723 | 21092 | 28829 | 15741 | 124275 |
| Protein residues | 11684 | 3069 | 12084 | 2537 | 3273 | 2183 | 2399 | 1302 | 7798 |
| Nucleotide residues | 2798 | 716 | 2887 | 694 | 715 | 165 | 439 | 252 | 2897 |
| **B Factors** (Å$^2$) | | | | | | | | | |
| Protein | 17.2 | 38.77 | 11.98 | 27.73 | 18.04 | 61.77 | 28.38 | 80.9 | 11.37 |
| Nucleotide | 24.61 | 51.56 | 21.91 | 44.42 | 26.67 | 93.74 | 36.22 | 133.64 | 18.66 |
| Ligand/ion | 28.87 | 44.84 | 10.67 | 38.69 | 31.5 | N/A | N/A | 113.67 | 23.73 |
| **RMSD** | | | | | | | | | |
| Bond lengths (Å) | 0.004 | 0.004 | 0.003 | 0.008 | 0.003 | 0.003 | 0.003 | 0.003 | 0.005 |
| Bond angles (°) | 0.603 | 0.587 | 0.552 | 0.679 | 0.443 | 0.549 | 0.58 | 0.673 | 0.683 |
| **Validation** | | | | | | | | | |
| MolProbity score | 1.7 | 1.59 | 1.68 | 1.64 | 1.54 | 1.76 | 1.53 | 2.1 | 1.78 |
| Clashscore | 7.29 | 6.23 | 6.98 | 5.97 | 5.01 | 7.53 | 6.36 | 13.19 | 8.74 |
| Rotamer outliers (%) | 0.01 | 0.04 | 0.03 | 0.05 | 0 | 0.1 | 0 | 0 | 0.09 |
| **Ramachandran plot** | | | | | | | | | |
| Favored (%) | 95.64 | 96.29 | 95.78 | 95.44 | 95.94 | 94.99 | 96.96 | 92.55 | 95.51 |
| Allowed (%) | 4.3 | 3.71 | 4.18 | 4.48 | 4.06 | 5.01 | 2.96 | 7.29 | 4.46 |
| Outliers (%) | 0.06 | 0 | 0.04 | 0.08 | 0 | 0 | 0.09 | 0.16 | 0.03 |
